# Enhanced neural tracking of the fundamental frequency of the voice

**DOI:** 10.1101/2020.10.28.359034

**Authors:** Jana Van Canneyt, Jan Wouters, Tom Francart

## Abstract

**Objective:** ‘F0 tracking’ is a novel method that investigates the neural processing of the fundamental frequency of the voice (f0) in continuous speech. Through linear modelling, a feature that reflects the stimulus f0 is predicted from the EEG data. Then, the neural response strength is evaluated through the correlation between the predicted and actual f0 feature. The aim of this study was to improve upon this ‘f0 tracking’ method by optimizing the f0 feature.

**Approach:** Specifically, we aimed to design a feature that approximates the expected EEG responses to the f0. We hypothesized that this would improve neural tracking results, because the more similar the feature and the neural response are, the easier it will be to reconstruct the one from the other. Two techniques were explored: a phenomenological model to simulate neural processing in the auditory periphery and a low-pass filter to approximate the effect of more central processing on the f0 response. Since these optimizations target different aspects of the auditory system, they were also applied in a cumulative fashion.

**Results:** Results obtained from EEG evoked by a Flemish story in 34 subjects indicated that both the use of the auditory model and the addition of the low-pass filter significantly improved the correlations between the actual and reconstructed feature. The combination of both strategies almost doubled the mean correlation over subjects, from 0.78 to 0.13. Moreover, canonical correlation analysis with the modelled feature revealed two distinct processes contributing to the f0 response: one driven by the compound activity of auditory nerve fibers with center frequency up to 8 kHz and one driven predominantly by the auditory nerve fibers with center frequency below 1 kHz.

**Significance:** The optimized f0 features developed in this study enhance the analysis of f0-tracking responses and facilitate future research and applications.

## 1. Introduction

Traditionally, auditory-evoked potentials are evoked by short repetitive stimuli, but research is progressing towards the use of continuous speech stimuli. Experiments with these natural stimuli are more pleasant for subjects and yield detailed information on auditory processing in day-to-day communication (Hamilton and Huth, 2020). As part of this movement, researchers developed a framework to analyse neural responses to continuous speech based on linear decoding models (e.g. Mesgarani et al. (2009); Lalor and Foxe (2010); Ding and Simon (2012); Crosse et al. (2016); Vanthornhout et al. (2018)). A linear decoding model, or backward model, reconstructs a specific stimulus-related feature from a linear combination of multi-channel neural responses and their time-lagged versions (Mesgarani et al., 2009). These linear models can be constructed for various stimulus features and depending on the feature, different aspects of auditory processing can be targetted. In this study, we focus on brainstem-dominated responses to the fundamental frequency of the voice (f0) in continuous speech, or “f0-tracking”, as described in Etard et al. (2019) and Van Canneyt et al. (2020b). Specifically, we aimed to optimize the feature that is used in these paradigms.

The performance of backward decoding models is evaluated based on the correlation between the reconstructed feature, derived from the EEG (or MEG), and the actual feature, derived from the stimulus. For f0-tracking, the actual feature is typically obtained by band-pass filtering the stimulus (or through empirical mode decomposition (Forte et al., 2017)). However, the EEG response is not a perfect reflection of the stimulus and therefore the EEG-derived feature and these stimulus-derived features cannot be expected to correlate perfectly. The EEG response is shaped by neural processes like adaptation, saturation, and refractory periods, which have been extensively studied and can be simulated with models of the auditory system. Moreover, researchers have studied the EEG response and its dependency on the evoking stimulus and defined important temporal and spectral response characteristics. The goal of this study was to use the available knowledge on phase-locked EEG responses to adjust the feature used for f0 tracking, such that it is more similar to what is expected from the EEG response. We hypothesized that this would improve the correlations obtained with linear modelling, as it would be easier to predict the feature from the EEG responses. Typically, correlations for f0-tracking responses are quite small, i.e. in the range of 0.03-0.08, so increasing these values is desired.

Two strategies were set out to optimize the f0 feature. In a first step, we aimed to account for a series of neural processes occurring in the auditory periphery. This included frequency-specific basilar membrane delays, adaptation effects and refractory effects in the primary auditory nerve fibers (ANF). For this purpose, we employed a phenomenological model of the auditory periphery (Carney, 1993; Zhang et al., 2001; Bruce et al., 2003; Zilany and Bruce, 2006, 2007; Zilany et al., 2009, 2014; Bruce et al., 2018). The model predicts neural firing patterns in a large population of ANFs based on the input stimulus. By summing together the firing patterns over the ANFs, the response on the population level can be estimated. In a previous study, this population response has been found to accurately simulate phase-locked responses to stimulus envelope modulations (Van Canneyt et al., 2019). Since the f0 manifests as envelope modulations, we expect the simulations to approximate the neural response to the f0 in continuous speech as well. Therefore, we hypothesized that using the simulated population response as a feature would increase the performance of the linear models.

An additional benefit of using the model of the auditory periphery by Bruce et al. (2018) is that relative contributions of neural populations with different center frequencies (CF) could be investigated. The adult human f0 ranges from about 80 to 300 Hz, and intuitively one would expect the f0 response to be driven by ANF with a CF in this range. However, there is evidence (from classic envelope following response (EFR) paradigms) that the f0 response is not primarily driven by the stimulus f0 but mostly by its harmonics (Aiken and Picton, 2006; Laroche et al., 2013), whose combined response periodicity equals the f0. For this reason, amongst others, ANF with larger CF are thought to contribute as well (Dau, 2003). Moreover, the higher harmonics of a stimulus can be divided in resolved and unresolved harmonics (Micheyl and Oxenham, 2004). Resolved harmonics are low frequency harmonics (< 1 kHz) which are each processed in a separate auditory filter in the cochlea. In contrast, unresolved harmonics have higher frequencies and multiples of them will occur within a single auditory filter so the auditory system processes them in a combined fashion. Several studies have tried to distinguish the contributions of resolved and unresolved harmonics to the classic EFR (Krishnan and Plack, 2011; Laroche et al., 2011, 2013), with varying results. Recently, findings of Saiz-Alia and Reichenbach (2020) suggested that fibers with CFs up to 8 kHz (corresponding to both resolved and unresolved harmonics) contribute more or less equally to the continuous f0-tracking response, but the stimulus used in that study has unnaturally strong higher harmonics (see discussion in Van Canneyt et al. (2020b)). We used the model simulations and canonical correlation analysis (CCA) to verify this finding for speech with a more natural speech profile.

With the model of the auditory periphery, EEG response characteristics up to primary auditory nerve are adequately captured. However, the f0 tracking response is predominantly generated beyond the ANFs. In our previous work, Van Canneyt et al. (2020b), we have shown that the primary sources for the f0 tracking response are located in the brainstem, with possible cortical contributions. Therefore, the second strategy focussed on auditory processing higher-up the auditory pathway. Auditory models of brainstem processing already exist (Nelson and Carney, 2004; Verhulst et al., 2018; Carney et al., 2015; Saiz-Alia and Reichenbach, 2020), but we chose to design a new model that is simple, yet highly effective for our purpose, by focussing on the spectrum of the response. It is known that the frequency limit for phase-locking decreases along the auditory pathway, causing cortical sources to contribute more strongly for stimuli with low f0.

This ties together with the fact that f0 (or envelope) following responses decrease in strength with increasing stimulus f0 (e.g. Purcell et al., 2004; Gransier, 2018; Van Canneyt et al., 2020a,b). The exact relation between response amplitude and stimulus frequency varies widely across individuals, and there are many peaks and valleys (Tichko and Skoe, 2017), but we hypothesized that this frequency-amplitude relation could be approximated with a Butterworth low-pass filter. Therefore, our higher-level model is essentially a low-pass filter for which we optimized the filter parameters, i.e. order and frequency cut-off, in a data-driven way. We hypothesized that applying this filter to the feature would enhance the backward modelling correlations because the spectrum of the EEG and the to-be-predicted feature match more closely.

In summary, this study aimed to optimize the feature used in linear models to analyse neural f0-tracking by incorporating prior knowledge of the f0 response. Two strategies were examined: 1) using simulations of the neural population response in the auditory periphery as the feature and 2) applying a low-pass filter to the feature to account for the effect of more central processing on the spectrum of the response. The two strategies were applied separately as well as combined, and the effect on backward modelling correlations was investigated. Additionally, the model simulations were used to quantify the relative contributions of ANF with different CF to the f0 response.

## 2. Methods

### 2.1. Dataset

The neural responses analysed in this study are part of an existing data set (Accou et al., 2020; Monesi et al., 2020) that was also used in our previous work (Van Canneyt et al., 2020b). EEG responses to continuous speech were measured for 34 young normal hearing participants, who were native Flemish (or Dutch) speakers (31 females, 3 males), with ages ranging between 18 and 24 years old (mean = 22.4 years, standard deviation = 1.4 years). All participants were normal hearing (all thresholds < 20 dB HL), which was verified using pure-tone audiometry (octave frequencies between 125 and 8000 Hz). The continuous speech stimulus was a Flemish story, titled “Milan” (written and narrated by Stijn Vranken), which lasted 14.6 minutes and had a mean f0 of 107 Hz (interquartile range = 34.7 Hz). The experiments were approved by the medical ethics committee of the University Hospital of Leuven and all subjects signed an informed consent form before participating (s57102).

### 2.2. EEG responses

The EEG responses in the dataset were recorded with a 64-channel Biosemi ActiveTwo EEG recording system (fs = 8192 Hz). The 64 Ag/AgCl active scalp electrodes were placed on a cap according to the international standardized 10-10 system (American Clinical Neurophysiology Society, 2006). Subjects were seated in an electromagnetically-shielded sound-proof booth and instructed to listen carefully to the story, which was presented binaurally through electrically-shielded insert phones (Etymotic ER-3A, Etymotic Research, Inc., IL, USA) using the APEX 3 software platform (Francart et al., 2008). Stimulus intensity was set to 62 dB A in each ear. The setup was calibrated in a 2-cm^3^ coupler (Brüel & Kjaer, type 4152, Nærum, Denmark) using stationary speech weighted noise with the same spectrum as the story. To encourage attentive listening, participants answered a question about the content of the story after its presentation.

We applied several preprocessing steps to the raw EEG data from the dataset. First, the data was downsampled to a sampling frequency of 1024 Hz. Then, artefacts were removed using a multi-channel Wiener filter algorithm with delays from −3 to 3 samples included and a noise weighting factor of 1 (Somers et al., 2018). The data was re-referenced to the average of all electrodes and band-pass filtered with a Chebyshev filter with 80 dB attenuation at 10 % outside the pass-band and a pass-band ripple of 1 dB. The filter cut-offs, i.e. 75 and 175 Hz, were chosen based on the f0 distribution of the story. We also applied a notch filter to remove the artefact caused by the third harmonic of the utility frequency at 150 Hz (the other affected frequencies did not fall in the bandpass filter range). The EEG was normalized to be zero mean with unit variance.

### 2.3. Linear decoding model

The EEG responses were analysed with linear backward decoding models implemented in MATLAB R2016b (The MathWorks Inc., 2016) using custom scripts and the mTRF toolbox (Crosse et al., 2016). A description of the main methods is provided here, but for details we refer to Van Canneyt et al. (2020b). In backward linear modelling or decoding, one reconstructs a known stimulus-related feature based on a linear combination of the time-shifted data from the EEG electrodes. In this study, time shifts between 0-40 ms in steps of 1/fs (fs = 1024 Hz) were included. First, a section of the data (including minimum 2 minutes of voiced data) was set aside for testing and the model was estimated based on the remainder of the data. Regularization was done using ridge regression (Tikhonov and Arsenin, 1977; Hastie et al., 2001; Machens et al., 2004). Then, the estimated model was used to reconstruct the feature for the testing data. Finally, the bootstrapped Spearman correlation between the reconstructed feature and the actual f0 feature, for the test section, was calculated (median over 100 index-shuffles). Importantly, unvoiced and silent sections were removed from the reconstructed and actual feature before correlating, because they have no reliable f0 (Forte et al., 2017). To validate the backward decoding results, we used a 3-fold cross-validation approach. The final backward correlation, i.e. the median over the folds, was compared to a significance level (based on correlations with spectrally-matched noise signals) to evaluate its statistical significance (two-sided test, *α* = 0.05).

### 2.4. The features

To investigate how the neural system tracks the f0, the linear modelling approach requires a f0 feature, i.e. a waveform reflecting the instantaneous f0 of the stimulus. In previous f0-tracking work (Van Canneyt et al., 2020b; Etard et al., 2019), f0 features were obtained by bandpass filtering the stimulus (’default’). The aim of this study was to develop a more optimal method to create the feature. A first strategy we explored, was to use a model of the auditory periphery to obtain the feature (’model’). A second strategy was to apply an additional low pass filter, that roughly simulates neural processing beyond the auditory periphery, either to the default feature (default + low-pass) or to the model feature (’model + low-pass’). The four features (the default feature, the model feature, the low-passed default feature and the low-passed model feature) are visualised in figure 1. Below the calculation of each of the features is discussed in detail. Importantly, unvoiced and silent sections were set to zero in all features before normalizing to zero mean and variance of 1. We performed linear decoding analysis of the data with each of the four features and compared the resulting correlations. Feature-induced differences between backward correlations were statistically evaluated in R (version 3.6.3., R Core Team (2018)), using linear mixed models (package lme4, version 1.1.21, Douglas et al. (2015)) with a random intercept per subject.

**Figure 1:**
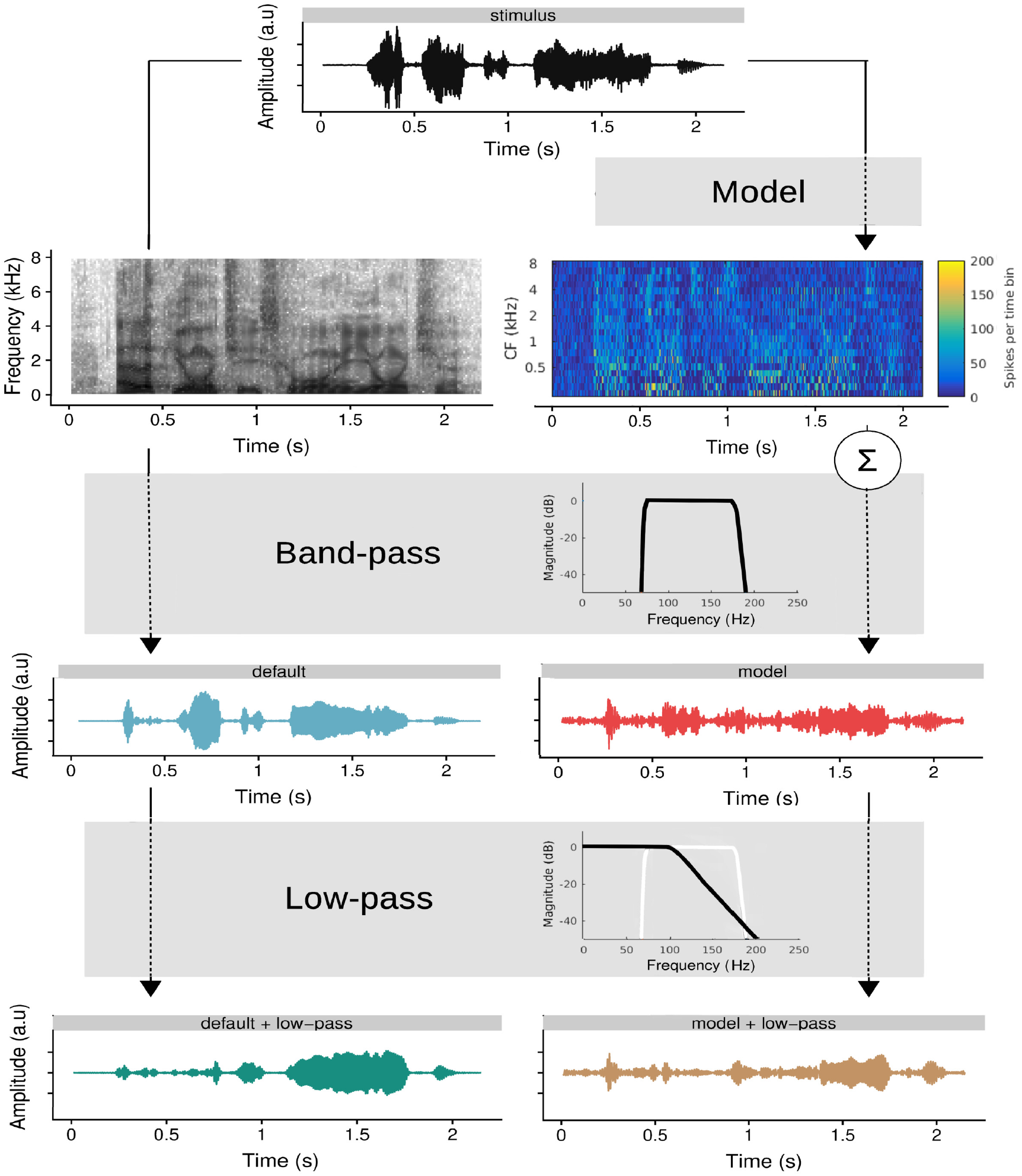
Visualisation of the stimulus and how the features were derived from it. The sentence shown is “Elk jongetje is gewoon een jongetje” (translation: “every boy is just a boy”).

#### 2.4.1. The default feature

The “default” feature was based on band-pass filtering of the stimulus. Specifically, we used a Chebyshev bandpass filter with 80 dB attenuation at 10 % outside the pass-band and a pass-band ripple of 1 dB. The filter cut-offs, i.e. 75 and 175 Hz, were chosen based on the f0 distribution of the story. This filter is identical to the one applied to the EEG (see above). The amplitude response of the band-pass filter, as well as the resulting default feature, is visualized in Figure 1.

#### 2.4.2. The model-based feature

The model-based feature is generated with a phenomenological model of the auditory periphery (Carney, 1993; Zhang et al., 2001; Bruce et al., 2003; Zilany and Bruce, 2006, 2007; Zilany et al., 2009, 2014; Bruce et al., 2018). The model simulates spike patterns from a population of auditory nerve fibers in response to an input stimulus. Here, the model simulated 20 different center frequencies (CFs) logarithmically spaced between 250 and 8000 Hz and for every CF, there were 50 nerve fibers with different spontaneous firing rates: 10 low (0.1 spikes/s), 10 mid (4 spikes/s) and 30 high (70 spikes/s). For a detailed description of the model as well as the model code, we refer to Bruce et al. (2018). However, two important changes were made to the model to increase the temporal resolution of the output: the window length of the smoothing Hamming window in the post-stimulus time-histogram (PSTH) was decreased from 128 to 32 samples and the amount of bins over which the PSTH was integrated was decreased from 10 to 5. The process to obtain the model-based f0 feature is visually represented in figure 1: the model received the Flemish story as input and produced simulated spike patterns for ANF at each of the CF, which can be visualised in a neurogram. The spike patterns were summed across all CFs (i.e. summing along the y-axis of the neurogram) to obtain the neural response at population level. Finally, the same band-pass filter as discussed in section 2.4.1 was applied to extract the neural response to the f0.

#### 2.4.3. The low-pass filtered features

We applied a low-pass filter to the feature such that the spectrum of the feature better resembled the spectrum of the expected f0 response, i.e. with reduced amplitude for higher frequencies. To avoid unwanted side effects of the filtering, especially in the stopband, we used a Butterworth filter. The order and cut-off frequency were determined in a data-driven way: for each subject, we calculated linear decoding models based on the default feature, low-pass filtered with different filter orders (1, 2, 4, 6, 8, 10, 12) and filter cut-offs (75, 80, 90, 100, 110, 120, 130, 140, 155, 175 Hz). Including a wider range of cut-offs made little sense because the features are already filtered by a bandpass filter that strongly attenuated frequencies outside this range (see earlier). The results of this optimisation are discussed in detail in section 3.2. In summary, we found a 8th (or higher) order filter with a cut-off frequency of 110 Hz to be optimal. The amplitude response of this filter is shown in figure 1. The same optimization process was performed for the model-based feature leading to nearly identical results, which were therefore not reported. As shown in figure 1, the optimized low-pass filter was applied to both the default and the model-based feature to create the two low-passed features.

### 2.5. The relative contribution of nerves with different center frequencies

The model simulations produced neural firing patterns for a group of 50 ANF at 20 CF, which were all summed together to obtain the model-based feature. In an additional analysis, we investigated the response at different CFs separately using a canonical correlation analysis (CCA). In preparation for the CCA, the spike patterns at each of the CF were filtered with the same bandpass filter specified earlier in section 2.4.1 and normalized to be zero mean. Moreover, the silent and unvoiced section were removed. Whereas linear backward decoding models are trained by finding the weighted combination of EEG channels that maximally correlates with a fixed feature, canonical correlation analysis (CCA) optimizes the correlation by applying weights to both the EEG channels and a set of features. In this case, the CCA assigned weights to the simulated response at each of the CFs, which is indicative of the relative importance of nerves with that CF for the f0 response. The CCA also determined weights for the EEG channels and their time-shifted versions (0-40 ms with 1/fs steps (fs = 1024 Hz)), but interpreting these ‘backward’ weights as a spatial distribution of the response is not reliable. As argued by Haufe et al. (2014a), large weights may be paired with channels unrelated to the signal of interest while channels containing response energy may receive small weights. These misleading effects occur because the linear model attempts to suppress noise components. To resolve this issue, Haufe et al. (2014a) proposed to transform backward models into forward models. In forward modelling, the EEG data in each recording channel is predicted based on the feature and its time-shifted versions. This method is less powerful than backward modelling, but since each EEG channel is treated separately, noise suppression cannot take place so the forward modelling weights can be reliably interpreted.

CCA estimated as many canonical components (sets of weights) as there are elements in the smallest set, which in this case was determined by the amount of CFs included in the model, i.e. 20. Each of these components was estimated under the constraint that they are uncorrelated with the previous components. The 20 resulting models, or CCA components, were applied to 2 minutes of unseen voiced data and bootstrapped Spearman correlations between the reconstructed features and the actual f0 features were calculated (median over 100 index-shuffles). To assess the significance of each of the components, significance thresholds were estimated in the same way as for the linear decoding models.

To understand the spatio-temporal characteristics of the canonical components, the significant components were transformed to a forward model, following Haufe et al. (2014a). This was done by weighing the model simulated responses at different CFs according to the weights estimated by CCA (instead of equal weighting in the default case) and summing it together to obtain a new f0 feature. This new feature and its time-shifted versions (−20 to 80 ms with 1/fs steps (fs = 1024 Hz)) were then used to predict the EEG response in each channel. The weights of the forward model can be interpreted through temporal response functions (an average over channels in function of time), which reflect the impulse response of the auditory system, and also through topoplots, which reveal the spatial distribution of the response at specific time lag. Because of the large degree of autocorrelation present in the f0 feature, response energy is spread in time, both in the TRFs and the topoplots. To help with interpretation, we calculated Hilbert TRFs, but the underlying autocorrelative smearing should be kept in mind. For more details on Hilbert TRFs and other aspects of the forward modelling, we refer to our previous work: Van Canneyt et al. (2020b).

## 3. Results

### 3.1. Comparison of backward decoding results

We performed linear decoding analysis of the same neural data with four different features: the default feature, the model-based feature, the low-passed default feature and the low-passed model-based feature. Figure 2 compares the backward correlations obtained for all subjects with each of the features. Visual comparison indicates that analysis with the model based-feature produced larger correlations compared to analysis with the default feature. Moreover, adding the low-pass filter improved correlations both for the default and the model-based feature. Significance levels are highly similar across features (dashed lines). The only feature that provided significant correlations for all subjects is the low-passed model-based feature. A linear mixed model with random intercept per subject was used to statistically evaluate the relative performance of the features. There was a significant difference between the correlations obtained with the default and the model-based feature (*β* = 0.030, df = 102, t = 10.7, p < 0.001). Moreover, there was a significant difference between the correlations obtained with the default and low-passed feature (*β* = 0.036, df = 102, t = 12.8, p < 0.001). Finally, the combination of the low-pass filter and the model-based feature resulted in significantly different correlations compared to the model-based feature on its own (*β* = 0.022, df = 102, t = 8.0, p < 0.001).

**Figure 2:**
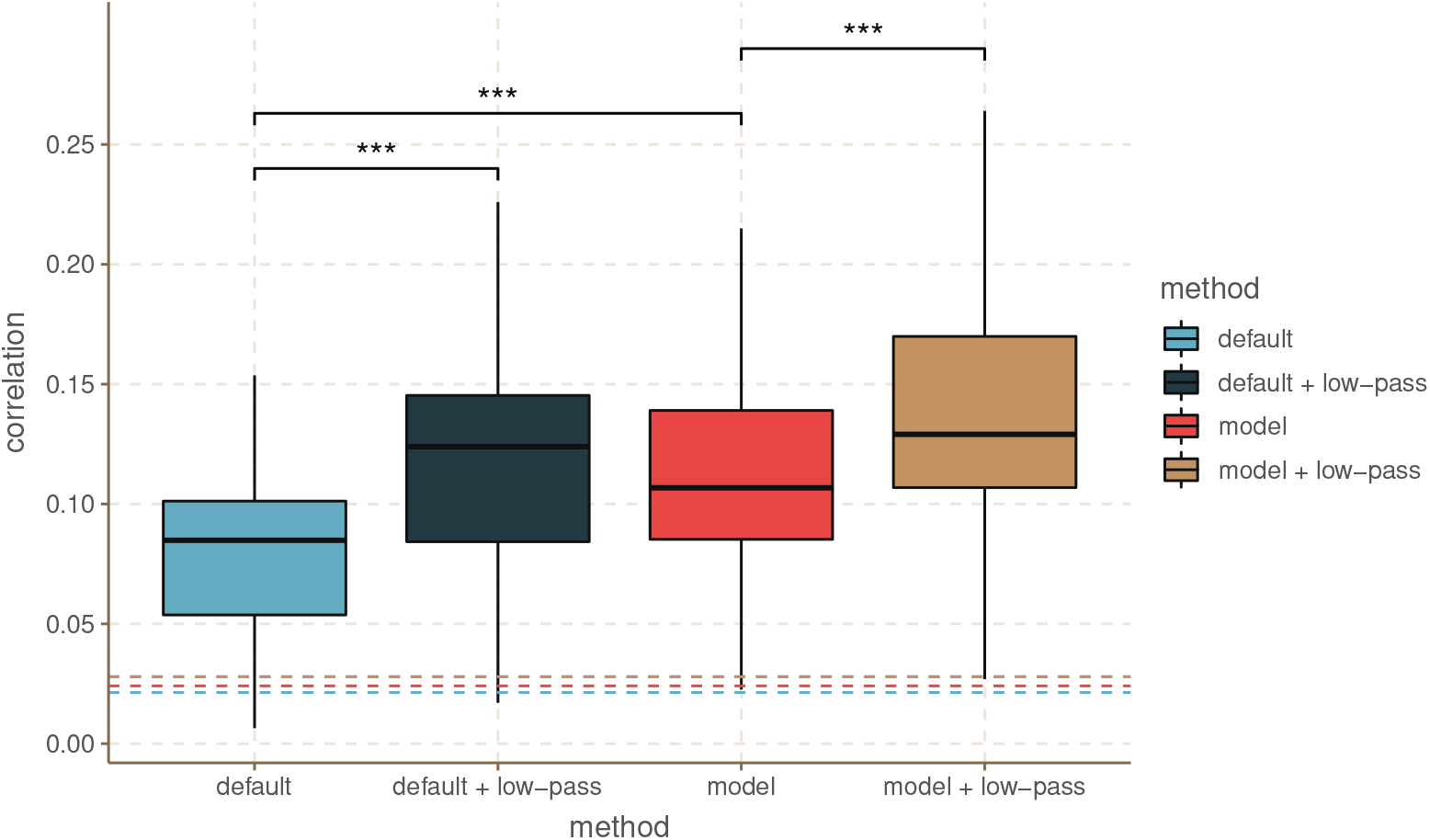
Comparison of correlations for all subjects obtained with each of the features. The dashed lines indicate the significance level. *** indicates a significant difference with a p < 0.001.

### 3.2. Optimisation of the low-pass filter

As described in the methods, the parameters of the Butterworth filter, used to filter the features, were defined in a data driven way. In figure 3, the results of this optimization are presented. We identified the filter parameters that induced the largest increase in correlation, compared to the correlation obtained with the default non-filtered feature. The results indicated that the largest increase in correlations, on average over subjects, occurred for a filter of 8th (or higher) order with a cut-off frequency of 110 Hz (panel A). For the majority of the subjects, increasing the order of the filter up to 8, while keeping the cut-off frequency fixed at 110 Hz, resulted in a monotonic increase of the correlation. Using filter orders larger than 8 did not further enhance the correlations (Panel B). With a fixed filter order of 8, a cut-off frequency of 110 Hz was most optimal for the majority of the subjects (n = 19), but for some subjects a cut-off of 100 Hz (n = 9), 120 (n = 5) or 130 Hz (n = 1) was better (Panel C). For filter cut-off frequencies near 175 Hz, the change in correlation induced by low-pass filtering approached 0, because in those cases the attenuation of the low-pass filter fell outside the bandpass-filter (applied earlier), and therefore had no effect. In contrast, filter cut-offs below 80 Hz tended to decrease the correlation, indicating the importance of the lower frequencies. Optimisation of the low-pass filter on the model-based feature led to highly similar results and was therefore not shown.

**Figure 3:**
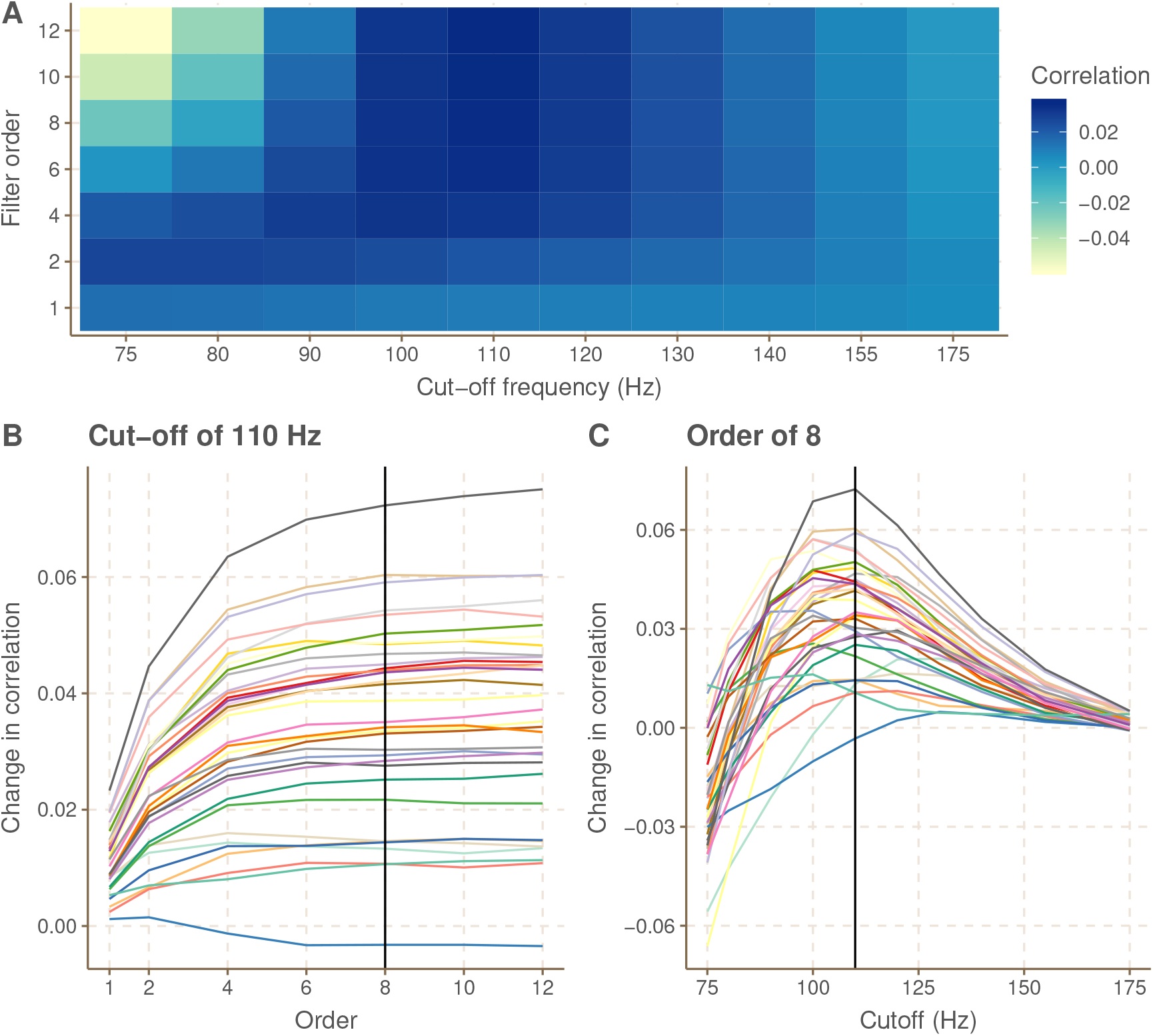
Results of the optimisation of the low-pass filter applied to the default feature A. Change in backward correlation caused by applying a low-pass filter with the specified order and cut-off frequency to the default feature, averaged over subjects. B. Change in backward correlation by altering the filter order with the cut-off frequency fixed at 110 Hz, for each subject separately. C. Change in backward correlation by altering the filter cut-off frequency with the filter order fixed at 8, for each subject separately.

### 3.3. The relative contribution of nerves with different center frequencies

To estimate the relative contribution of auditory nerve fibers with different CF to the f0 response, we performed CCA with the simulated spike patterns per CF. Out of the 20 estimated CCA components, the first two provided correlations that were larger than the significance level for the majority of the subjects (Figure 4, panel A). The median correlation over subjects obtained for the first component (0.099) is similar to what was found with the model-based feature in regular linear decoding (0.106), while the median backward correlation of the second component is smaller (0.076). The variance over subjects is also similar to what was observed for regular linear modelling. Panel B and C of figure 4 indicate the weight pattern for the first and second component, respectively. Note that the sign of these weights can be reversed without a change in meaning, as long as it is done for all the weights. The estimated weight patterns are highly similar across subjects. The first component revealed positive weights to all CFs except the lowest one, i.e. 250 Hz, which had a large negative weight. The second component is divided between positive weights for CFs below 1 kHz and smaller, (mostly) negative weights above 1 kHz. Weight patterns for non-significant CCA components were not analysed.

**Figure 4:**
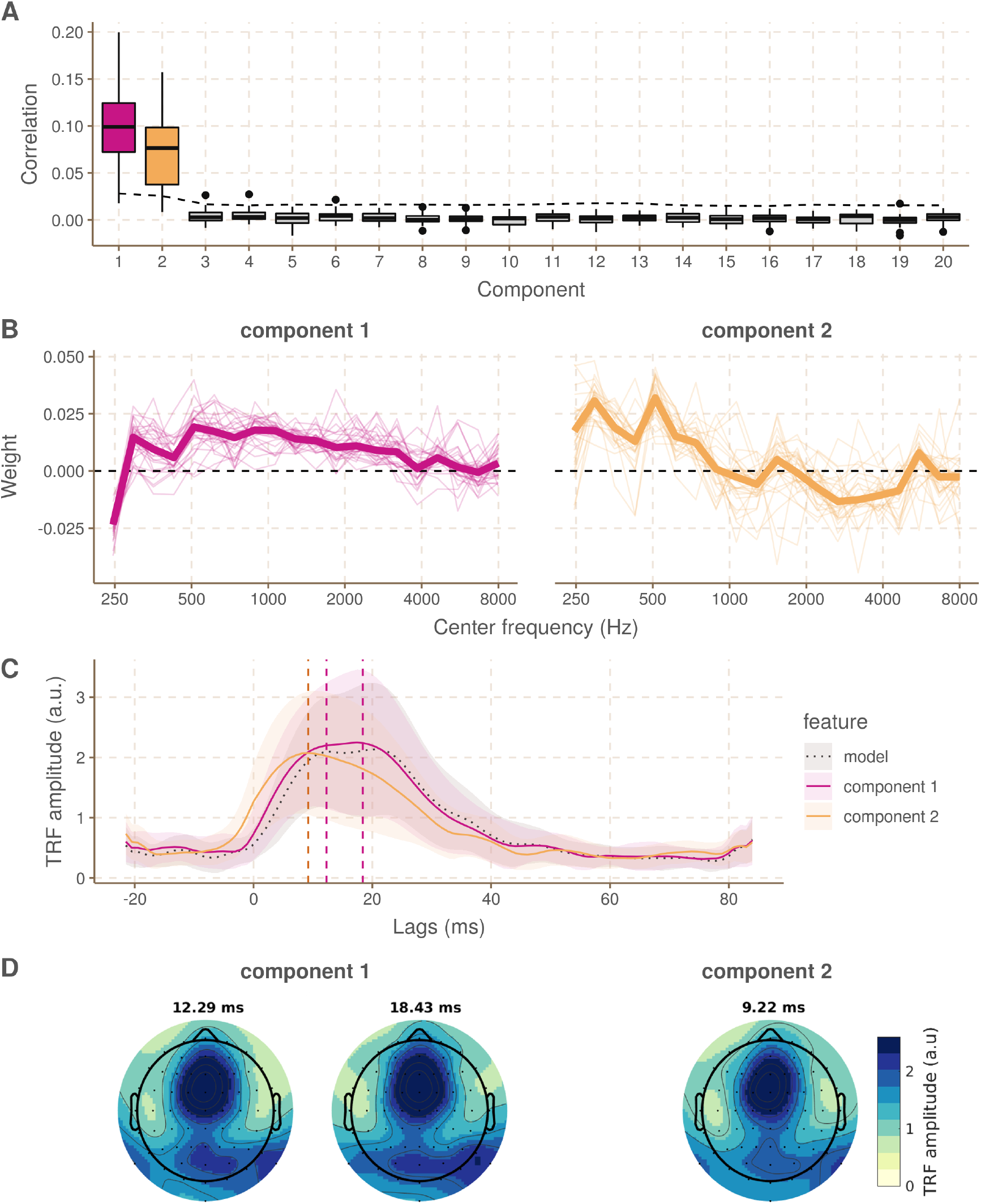
Results of the CCA using the spike patterns per CF and the EEG (+ time shifted versions) A. Backward correlations for each of the subjects and for each of the 20 canonical components. The significance level is indicated with a dashed line. B. CCA weights across CFs for the first and second component respectively, for each of the subjects (thin line) and in the median case (thick line). C. Hilbert TRFs for the two significant canonical components and the regular model (black dotted line). The peaks lags at which topoplots were plotted in panel D are indicated with vertical dashed lines. D. Topoplots at the peaks lags of the TRFs in panel C.

Through forward modelling using features assembled from the neurogram according to the weightings displayed in panel B of Figure 4, the spatio-temporal characteristics of the canonical components was analysed. Panel C of Figure 4 presents the Hilbert TRFs for the two significant canonical components. The TRF for the first component peaks around 12.3 and 18.4 ms and is highly similar to the TRF of the regular model feature (black dotted line). This is not suprising as the CCA weights approximate equal weighting across CFs. However, the second component has a more narrow and earlier peak at 9.22 ms. The topoplots in panel D of Figure 4 indicate the spatial distribution of the response energy at these peak lags and these are highly similar to what was reported in Van Canneyt et al. (2020b). The second component seems to have less temporo-mastoidal activity which, together with the narrower and earlier TRF, indicates less cortical contributions compared to component 1.

## 4. Discussion

The goal of this study was to enhance the analysis of f0-tracking responses to continuous speech by optimizing the feature used in the linear decoding models. Backward correlations for f0-tracking responses reported in earlier studies are typically quite small, i.e. in the range of 0.03 −0.08. Larger correlations would facilitate the detection and interpretation of group differences (less floor effects) and make f0-tracking analysis more robust. We hypothesized that better results would be obtained when the feature better resembled the expected neural response, as predicting the one from the other would be easier.

A first strategy to optimize the feature was to use a model of the auditory periphery to simulate the neural response to the stimulus at the level of the primary auditory nerve. In a prior study, Van Canneyt et al. (2019), we showed how simulated population responses constructed through this model reliably predict neural responses to envelope modulations. Here, the simulated population responses were used as a feature in the linear decoding models. The model-based feature improved the mean correlation over subjects from 0.079 to 0.109, compared to the default feature. The model simulated auditory processing up to the primary auditory nerve, but f0-tracking is generated in the brain stem, with possible cortical contributions (Van Canneyt et al., 2020b). To account for the higher processing stages, we focussed on simulating the limitations of phase-locking. Phase-locking is less reliable for higher frequencies and the higher up the auditory pathway, the lower the maximum frequency that can be phase-locked to. This leads to a decreasing amplitude-frequency relation for the neural response, which we simulated through low-pass filtering. As shown in figure 2, low-pass filtering the default feature improved the mean correlation over subjects from 0.079 to 0.115. Since the two strategies target processes from different sections of the auditory pathway, it made sense to evaluated their combined effect. The combination of both strategies delivered the best results with significant correlations for all subject and almost a doubling of the mean correlation across subjects, from 0.079 to 0.130.

Importantly, the newly developed features differ in the time and computational resources necessary to obtain them. Depending on the duration of the continuous speech stimulus, calculating the simulated neural responses with the phenomenological model is computationally very expensive. In experimental settings where the same stimulus is presented to many subjects, use of the model is feasible as the model simulation can be reused for all subjects. However, the process to obtain the model-based feature is likely too slow for real-time applications. In contrast with the model-based feature, the addition of a low-pass filter is a quick and simple operation, which is easy to implement and likely possible in real-time. Moreover, even though this approach is relatively rudimentary, our results indicate it still provides a substantial benefit. Alternatively, one could account for auditory processing beyond the auditory nerve, by using a model of the auditory pathway up to the brainstem, as proposed by Verhulst et al. (2018) or Saiz-Alia and Reichenbach (2020). This way, neural responses at the level of the brainstem are simulated. However, these models are even more computationally expensive than the model of the auditory periphery.

The parameters of the filter used for the low-passed features were determined in a data-driven way. On a group-level, the backward correlations improved the most when the feature was filtered with a 8th (or higher) order Butterworth filter with a cut-off at 110 Hz. The order of the filter could be further increased without impacting the correlations but the optimal cut-off frequency was rather specific: varying it more than 10 Hz up or down reduced the correlations. It is also possible to use the optimal filter parameters for each subject individually, however this barely improved the correlations on a group-level (from 0.1145 to 0.1172 for the low-passed default feature and from 0.1309 to 0.1313 for the low-passed model-based feature). The optimization process was time-intensive and useful to develop the new feature, but does not necessarily need to be repeated for new data/stimuli. From explorations on different datasets with different evoking stimuli, we have learned that the optimal filter order is usually situated between 4 and 8, with voices with higher f0 favouring lower order filters. The optimal filter cut-off usually falls a little (e.g. 40-50 Hz) above the lower cut-off chosen for the bandpass filter, which is determined based on the f0 distribution of the story. Essentially, the filter should be designed such that the frequencies in the lower range of the f0 distribution of the stimulus are left untouched and higher frequencies are gradually more attenuated.

This study also included an investigation of the relative contributions of ANF with different CFs to the neural f0 tracking response. In this analysis, the simulated responses at different CF were assigned weights to optimize the correlation with a linear combination of the multi-channel and time-lagged EEG. The first CCA component indicated mainly positive weights, which confirms the findings by Saiz-Alia and Reichenbach (2020) that the f0-tracking response is generated by a collective of neurons with CFs up to 8kHz. The backward correlations obtained for this first CCA component were highly similar to the correlations obtained for the regular model-based feature, which makes sense since the weight pattern strongly resembles the uniform weighting used in the regular model-based feature. The CCA weights do indicate a steady decrease in relative contribution towards larger CF, which contrast the finding of Saiz-Alia and Reichenbach (2020) where CF up to 8 kHz were considered to contribute equally. Potentially, this difference is related to the fact that the stimulus of Saiz-Alia and Reichenbach (2020) has stronger higher harmonics than the stimulus of the present study. The observation that nerves with higher CF contribute to the neural f0 tracking response, not just the ANF with CF near the f0, follows the results of Dau (2003). Moreover, it also is in line with previous findings that claim that the EFR/f0-response is driven by both resolved and unresolved harmonics of the stimulus, not just the f0 (Jeng et al., 2011; Laroche et al., 2011, 2013; Van Canneyt et al., 2020a). Finally, the fact that higher harmonics are important drivers of the response could partly explain why the model feature outperforms the default feature: the model takes the full stimulus spectrum as input and can process the relative strengths of the higher harmonics and estimate their contribution to the f0 response, whereas the default feature only takes the energy around the f0 into account.

Remarkably, the CCA brought up a second component with (smaller) significant correlations, which is per definition uncorrelated to the first. CCA components differ in the weights assigned to the CF, but also have different temporal-spatial patterns, i.e. the weighting of different EEG channels at different time-shifts. Therefore, a second significant component could indicate an additional neural process underlying the f0-tracking response, possibly with different neural generators. The weights for the second component are large and positive for ANF with lower CF (<1000 Hz) and smaller and mostly negative for higher CF. This pattern could indicate that the process behind the second component focusses on the resolved harmonics in the stimulus and disregards the unresolved harmonics which typically occur above 1000 Hz. To learn more about the neural origin of this second response component, and how it differs from the first component, we applied Haufe et al. (2014b)’s suggestion to turn a backward model into a forward model. The results for the first component are highly similar to what was found for the regular model feature and to what was reported in our previous work (Van Canneyt et al., 2020b): TRFs with two peaks at lags around 13 and 18 ms and a topoplot with central and right temporo-mastoidal activity, suggestive of generators in the brainstem and right auditory cortex. The second process has a similar predominantly central spatial pattern but reduced tempero-mastoidal activity as well as only one and earlier TRF peak around 9 ms. This suggests that this second process occurs predominantly in the brainstem, without cortical contributions. These findings seem in line with the theory put forward by Laroche et al. (2011, 2013) that resolved and unresolved harmonics are processed in different but interacting pathways that converge in the upper brainstem.

## 5. Conclusion

In summary, this study has enhanced neural f0-tracking by optimizing the f0 feature such that it better resembles the expected neural response. Our recommendations are as follows: when fast and flexible implementation is required, low-pass filtering the feature is a great tool to boost correlations. When the stimulus is fixed and heavy computations are possible, the model-based feature, combined with a low-pass filter is preferred. Finally, if one wants to increase precision at the cost of even more computational power, one should consider a more extensive model of the auditory system that includes the brainstem (and ideally the primary auditory cortex as well). Besides, model simulations combined with CCA indicated that f0-tracking might be generated by two uncorrelated processes of which the first dominant one is driven by ANF with a broad range of CFs (up to 8 kHz) and the second smaller one is driven mostly by ANF responding to unresolved harmonics (CFs below 1 kHz). Cortical contributions are larger for the first process compared to the second.

## Acknowledgments

Authors would like to thank Bernd Accou and Wendy Verheijen for collecting the dataset used in this study. They were assisted in data collection by Amelie Algoet, Jolien Smeulders, Lore Kerkhofs, Sara Peeters, Merel Dillen, Ilham Gamgami en Amber Verhoeven. We also would like to thank Simon Geirnaert for his advice, especially with regards to CCA. This research was funded by TBM-project LUISTER (T002216N) from the Research Foundation Flanders (FWO) and also jointly by Cochlear Ltd. and Flanders Innovation & Entrepreneurship (formerly IWT), project 50432. Additionally, this project has received funding from the European Research Council under the European Unions Horizon 2020 research and innovation programme (grant agreement No. 637424, ERC starting grant to Tom Francart). The first author, Jana Van Canneyt, is supported by a PhD grant for Strategic Basic research by the Research Foundation Flanders (FWO), project number 1S83618N. Finally, the research is carried out with support from a Wellcome Trust Collaborative Award in Science RG91976 to Dr. Bob Carlyon and Jan Wouters, and with support from Flanders Innovation & Entrepreneurship through the VLAIO research grant HBC.2019.2373 with Cochlear. There are no conflicts of interest, financial, or otherwise.

## Notes

### Competing Interest Statement

The authors have declared no competing interest.

## References

Accou, B., Monesi, M. J., Montoya, J., Van Hamme, H., and Francart, T. (2020). Modeling the relationship between acoustic stimulus and EEG with a dilated convolutional neural network. In 28th European Signal Processing Conference (EUSIPCO), Amsterdam, Netherlands (in press).

Aiken, S. J. and Picton, T. W. (2006). Envelope following responses to natural vowels. Audiology and Neurotology, 11(04):213–232.

American Clinical Neurophysiology Society (2006). Guideline 5: guidelines for standard electrode position nomenclature. Am. J. Electroneurodiagnostic Technol., 46:222–225.

Bruce, I. C., Erfani, Y., and Zilany, M. S. (2018). A phenomenological model of the synapse between the inner hair cell and auditory nerve: Implications of limited neurotransmitter release sites. Hearing Research, 360:40–54.

Bruce, I. C., Sachs, M. B., and Young, E. D. (2003). An auditory-periphery model of the effects of acoustic trauma on auditory nerve responses. The Journal of the Acoustical Society of America, 113(1):369–388.

Carney, L. H. (1993). A model for the responses of low-frequency auditory-nerve fibers in cat. The Journal of the Acoustical Society of America, 93(1):401–417.

Carney, L. H., Li, T., and McDonough, J. M. (2015). Speech coding in the brain: Representation of vowel formants by midbrain neurons tuned to sound fluctuations. eNeuro, 2(4).

Crosse, M. J., Di Liberto, G. M., Bednar, A., and Lalor, E. C. (2016). The multivariate temporal response function (mTRF) toolbox: A MATLAB toolbox for relating neural signals to continuous stimuli. Frontiers in Human Neuroscience, 10(NOV2016).

Dau, T. (2003). The importance of cochlear processing for the formation of auditory brainstem and frequency following responses. The Journal of the Acoustical Society of America, 113(2):936–950.

Ding, N. and Simon, J. Z. (2012). Neural coding of continuous speech in auditory cortex during monaural and dichotic listening. Journal of Neurophysiology, 107(1):78–89.

Douglas, B., Maechler, M., Bolker, B., and Walker, S. (2015). Fitting Linear Mixed-Effects Models Using lme4. Journal of Statistical Software, 67(1):1–48.

Etard, O., Kegler, M., Braiman, C., Forte, A. E., and Reichenbach, T. (2019). Decoding of selective attention to continuous speech from the human auditory brainstem response. NeuroImage, 200(May):1–11.

Forte, A. E., Etard, O., and Reichenbach, T. (2017). The human auditory brainstem response to running speech reveals a subcortical mechanism for selective attention. eLife, 6:1–13.

Francart, T., van Wieringen, A., and Wouters, J. (2008). APEX 3: a multi-purpose test platform for auditory psychophysical experiments. Journal of Neuroscience Methods, 172(2):283–293.

Gransier, R. (2018). Phase-locked neural activity as a biomarker for auditory functioning: from speech perception to cochlear implant fitting. PhD thesis, KU Leuven.

Hamilton, L. S. and Huth, A. G. (2020). The revolution will not be controlled: natural stimuli in speech neuroscience. Language, Cognition and Neuroscience, 35(5):573–582.

Hastie, T., Tibshirani, R., and Friedman, J. (2001). The Elements of Statistical Learning. Springer, New York.

Haufe, S., Meinecke, F., Görgen, K., Dähne, S., Haynes, J. D., Blankertz, B., and Bießmann, F. (2014a). On the interpretation of weight vectors of linear models in multivariate neuroimaging. NeuroImage, 87:96–110.

Haufe, S., Meinecke, F., Görgen, K., Dähne, S., Haynes, J.-D., Blankertz, B., and Bießmann, F. (2014b). On the interpretation of weight vectors of linear models in multivariate neuroimaging. NeuroImage, 87:96–110.

Jeng, F. C., Costilow, C. E., Stangherlin, D. P., and Lin, C. D. (2011). Relative power of harmonics in human frequency following responses associated with voice pitch in American and Chinese adults. Perceptual and Motor Skills, 113(1):67–86.

Krishnan, A. and Plack, C. J. (2011). Neural encoding in the human brainstem relevant to the pitch of complex tones. Hearing Research, 275(1-2):110–119.

Lalor, E. C. and Foxe, J. J. (2010). Neural responses to uninterrupted natural speech can be extracted with precise temporal resolution. European Journal of Neuroscience, 31(1):189–193.

Laroche, M., Dajani, H., and Marcoux, A. (2011). Contribution of resolved and unresolved harmonic regions to brainstem speech-evoked responses in quiet and in background noise. Audiology Research, 1(1S).

Laroche, M., Dajani, H. R., Prèvost, F., and Marcoux, A. M. (2013). Brainstem auditory responses to resolved and unresolved harmonics of a synthetic vowel in quiet and noise. Ear and Hearing, 34(1):63–74.

Machens, C. K., Wehr, M. S., and Zador, A. M. (2004). Linearity of Cortical Receptive Fields Measured with Natural Sounds. Journal of Neuroscience, 24(5):1089–1100.

Mesgarani, N., David, S. V., Fritz, J. B., and Shamma, S. A. (2009). Influence of context and behavior on stimulus reconstruction from neural activity in primary auditory cortex. Journal of Neurophysiology, 102(6):3329–3339.

Micheyl, C. and Oxenham, A. J. (2004). Sequential F0 comparisons between resolved and unresolved harmonics: No evidence for translation noise between two pitch mechanisms. The Journal of the Acoustical Society of America, 116(5):3038–3050.

Monesi, M. J., Accou, B., Montoya-Martinez, J., Francart, T., and Van hamme, H. (2020). An LSTM based architecture to relate speech stimulus to EEG. In ICASSP, IEEE International Conference on Acoustics, Speech and Signal Processing-Proceedings. IEEE.

Nelson, P. C. and Carney, L. H. (2004). A phenomenological model of peripheral and central neural responses to amplitude-modulated tones. The Journal of the Acoustical Society of America, 116(4):2173–2186.

Purcell, D. W., John, S. M., Schneider, B. A., and Picton, T. W. (2004). Human temporal auditory acuity as assessed by envelope following responses. The Journal of the Acoustical Society of America, 116(6):3581–3593.

R Core Team (2018). R: A Language and Environment for Statistical Computing. R Foundation for Statistical Computing, Vienna, Austria.

Saiz-Alia, M. and Reichenbach, T. (2020). Computational modeling of the auditory brainstem response to continuous speech. Journal of Neural Engineering, in press:0–31.

Somers, B., Francart, T., and Bertrand, A. (2018). A generic EEG artifact removal algorithm based on the multi-channel Wiener filter. Journal of Neural Engineering, 15(3).

The MathWorks Inc. (2016). MATLAB: R2016b. Natick, Massachusetts.

Tichko, P. and Skoe, E. (2017). Frequency-dependent fine structure in the frequency-following response: The byproduct of multiple generators. Hearing Research, 348:1–15.

Tikhonov, A. N. and Arsenin, V. Y. (1977). Solutions of ill-posed problems. Scripta series in mathematics. V. H. Winston & Sons, Washington.

Van Canneyt, J., Hofmann, M., Wouters, J., and Francart, T. (2019). The effect of stimulus envelope shape on the auditory steady-state response. Hearing research, 380:22–34.

Van Canneyt, J., Wouters, J., and Francart, T. (2020a). From modulated noise to natural speech: The effect of stimulus parameters on the envelope following response. Hearing Research, 393:107993.

Van Canneyt, J., Wouters, J., and Francart, T. (2020b). Neural tracking of the fundamental frequency of the voice: male voices preferred. bioRxiv.

Vanthornhout, J., Decruy, L., Wouters, J., Simon, J. Z., and Francart, T. (2018). Speech Intelligibility Predicted from Neural Entrainment of the Speech Envelope. JARO-Journal of the Association for Research in Otolaryngology, 19(2):181–191.

Verhulst, S., Alto`e, A., and Vasilkov, V. (2018). Computational modeling of the human auditory periphery: auditory-nerve responses, evoked potentials and hearing loss. Hearing Research, 360:55–75.

Zhang, X., Heinz, M. G., Bruce, I. C., and Carney, L. H. (2001). A phenomenological model for the responses of auditory-nerve fibers: I. Nonlinear tuning with compression and suppression. The Journal of the Acoustical Society of America, 109(2):648–670.

Zilany, M. S. A. and Bruce, I. C. (2006). Modeling auditory-nerve responses for high sound pressure levels in the normal and impaired auditory periphery. The Journal of the Acoustical Society of America, 120(3):1446–1466.

Zilany, M. S. A. and Bruce, I. C. (2007). Representation of the vowel /epsilon/ in normal and impaired auditory nerve fibers: model predictions of responses in cats. The Journal of the Acoustical Society of America, 122(1):402–417.

Zilany, M. S. A., Bruce, I. C., and Carney, L. H. (2014). Updated parameters and expanded simulation options for a model of the auditory periphery. Journal of the Acoustical Society of America, 135(1):283–286.

Zilany, M. S. A., Bruce, I. C., Nelson, P. C., and Carney, L. H. (2009). A phenomenological model of the synapse between the inner hair cell and auditory nerve: Long-term adaptation with power-law dynamics. The Journal of the Acoustical Society of America, 126(5):2390–2412.

